## Supplementary Material for "Sleep does not influence schema-facilitated motor memory consolidation"

*Table S1: Group differences in participant characteristics and assessments of sleep and vigilances for Experiment 1.*

| **A. Variable** | **t** | **p** | **Cohen's d** |
|---|---|---|---|
| Age | -1.29 | 0.20 | -0.366 |
| BAI score | 0.90 | 0.38 | 0.253 |
| BDI score | 0.21 | 0.83 | 0.060 |
| Handedness score | -0.26 | 0.80 | -0.073 |
| PSQI score | 0.67 | 0.51 | 0.188 |
| Daytime sleepiness score | -1.68 | 0.10 | -0.475 |
| Sleep duration, 3 nights prior to S2 | -0.48 | 0.63 | -0.136 |
| SMS duration | -0.54 | 0.59 | -0.153 |
| SMS quality | 0.20 | 0.84 | 0.058 |
| **B. SSS** | **F** | **p** | **Partial $\eta^2$** |
| Session | 0.47 | 0.50 | 0.010 |
| Session x Group | 0.83 | 0.37 | 0.017 |
| Group | 0.62 | 0.43 | 0.013 |
| **C. PVT** | **F** | **p** | **Partial $\eta^2$** |
| Session | 1.04 | 0.31 | 0.021 |
| Session x Group | 0.001 | 0.98 | <0.001 |
| Group | 0.27 | 0.60 | 0.006 |

Output of statistical analyses assessing group differences (Nap vs. No Nap) in participant characteristics, sleep quality and quantity prior to the experimental sessions as well as subjective (Stanford Sleepiness Scale (SSS); Hoddes, Dement, & Zarcone, 1972) and objective (Psychomotor Vigilance Task (PVT); Dinges & Powell, 1985) assessments of vigilance. Means and SDs are provided in Table 1 of the main text. Variables in section **A** were assessed with independent samples t-tests (df = 48 for all). SSS (section **B**) and PVT (**C**) were assessed with 2 (Session) by 2 (Group) ANOVAs (df = 1,48 for all effects). No significant Group, Session or Group by Session effects were revealed. BAI = Beck's anxiety inventory [3]; BDI = Beck's depression inventory [4]; PSQI = Pittsburgh Sleep Quality Index [5], SMS = St. Mary's sleep questionnaire [6]; S2= session 2.

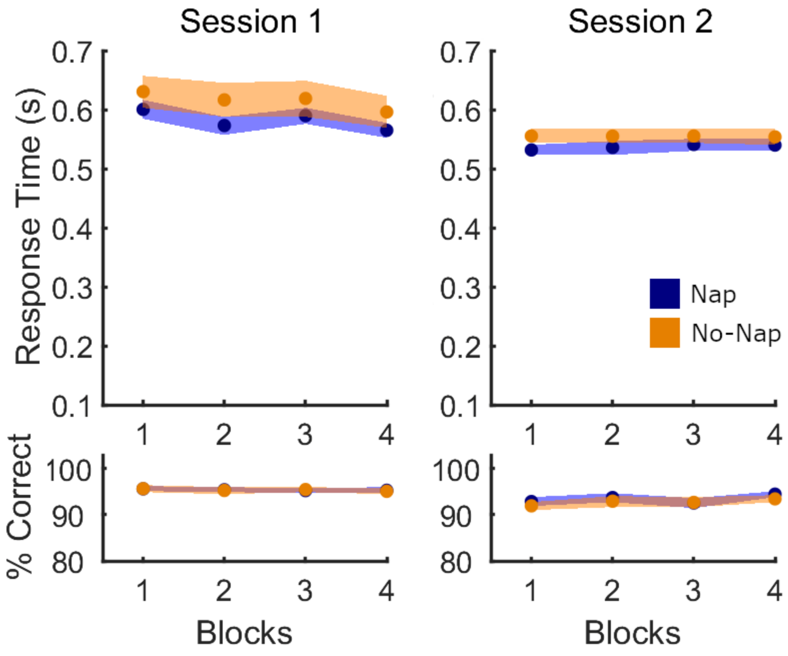


*Figure S1: Performance on the pseudo-random SRTT in Experiment 1. Mean response time (in seconds; top panel) and % correct transitions per block of task (bottom) are depicted separately for Session 1 and Session 2. Output of the corresponding statistical analyses is provided in Supplementary Table 2 below.*

*Table S2: Performance on the pseudo-random SRT task in Experiment 1.*

| Effect | df | F | p | Partial $\eta^2$ |
|---|---|---|---|---|
| **A. Response Time (RT)** | | | | |
| *Session 1* | | | | |
| Block | 2.5,121.9 | 11.59 | <0.001* | 0.195 |
| Block x Group | 2.5,121.9 | 0.59 | 0.60 | 0.012 |
| Group | 1,48 | 1.18 | 0.28 | 0.024 |
| *Session 2* | | | | |
| Block | 3,144 | 0.45 | 0.72 | 0.009 |
| Block x Group | 3,144 | 0.61 | 0.61 | 0.012 |
| Group | 1,48 | 1.40 | 0.24 | 0.028 |
| **B. Accuracy** | | | | |
| *Session 1* | | | | |
| Block | 3,144 | 0.26 | 0.86 | 0.005 |
| Block x Group | 3,144 | 0.06 | 0.98 | 0.001 |
| Group | 1,48 | 0.002 | 0.97 | <0.001 |
| *Session 2* | | | | |
| Block | 3,144 | 3.11 | 0.03 | 0.061 |
| Block x Group | 3,144 | 0.44 | 0.73 | 0.009 |
| Group | 1,48 | 0.30 | 0.59 | 0.006 |
| **C. Performance Index (PI)** | | | | |
| *Session 1* | | | | |
| Block | 3,144 | 5.50 | 0.001* | 0.103 |
| Block x Group | 3,144 | 0.56 | 0.64 | 0.012 |
| Group | 1,48 | 1.04 | 0.31 | 0.021 |
| *Session 2* | | | | |
| Block | 3,144 | 1.88 | 0.14 | 0.038 |
| Block x Group | 3,144 | 0.80 | 0.50 | 0.016 |
| Group | 1,48 | 1.27 | 0.27 | 0.026 |

Output of statistical analyses assessing group differences in performance on the pseudo-random SRT task assessing general motor execution administered prior to and following the sequential SRT task in Sessions 1 and 2, respectively. Separate 4 (Block) by 2 (Group) ANOVAs were run per each variable (**A**: Response Time, RT; **B**: Accuracy; and **C**: Performance Index, PI) and each Session. The presence of a significant effect of block during Session 1 for RT and PI indicates a general improvement in performance, presumably due to task familiarization (see Supplementary Figure 1, Session 1). No main effect of group or block x group interaction were observed for any measure or in either session, demonstrating that general motor execution did not differ between experimental groups. Df = degrees of freedom.

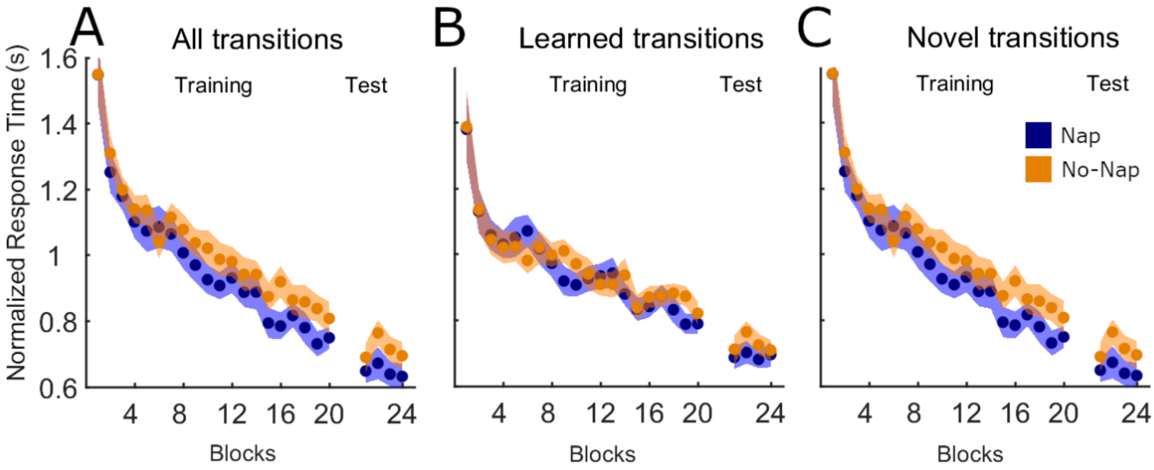


*Figure S2*: *Normalized response time during Session 2 training and test runs for all transitions (**A**), learned transitions (**B**), and novel transitions (**C**) for the two groups in Experiment 1. Shaded areas represent the SEM. Normalization was computed by dividing the response time per each block and participant by that individual's mean response time during the 4 blocks of Session 1 test. After normalization, the trend towards a significant group difference observed in the raw data (see main text, Figures 2 and 3) disappeared, as evidenced by the lack of group effects or group x block interaction. Detailed statistics are as follows:*

*All transitions – training – main effect group: $F(1,48)=0.65$, $p=0.42$, $\eta^2=0.013$; block × group interaction: $F(3.7,176.4)=0.59$, $p=0.66$, $\eta^2=0.012$; test – main effect group: $F(1,46)=2.09$, $p=0.16$, $\eta^2=0.043$; block × group interaction: $F(2.5,112.7)=0.31$, $p=0.78$, $\eta^2=0.007$.*

*Learned transitions – training –main effect group: $F(1,48)=0.09$, $p=0.77$, $\eta^2=0.002$; block × group interaction: $F(4.2,203.2)=0.75$, $p=0.57$, $\eta^2=0.015$; test – main effect group: $F(1,46)=1.14$, $p=0.29$, $\eta^2=0.024$; block × group interaction: $F(2.2,101.0)=0.25$, $p=0.73$, $\eta^2=0.008$.*

*Novel transitions – training –main effect group: $F(1,48)=1.39$, $p=0.25$, $\eta^2=0.028$; block × group interaction: $F(4.0,191.2)=0.47$, $p=0.76$, $\eta^2=0.010$; test – main effect group: $F(1,46)=1.87$, $p=0.18$, $\eta^2=0.039$; block × group interaction: $F(2.7,126.0)=0.13$, $p=0.93$, $\eta^2=0.003$.*

*Table S3: Performance on the generation task in Experiment 1.*

| Variable | Nap | No-Nap | t(48) | p | Cohen's d |
|---|---|---|---|---|---|
| *Session 1* | | | | | |
| % correct transitions | 66.1 (34.8) | 49.9 (35.2) | 1.63 | 0.11 | 0.462 |
| % correct ordinal positions | 56.8 (36.6) | 44.3 (37.5) | 1.19 | 0.24 | 0.337 |
| *Session 2* | | | | | |
| % correct transitions | 82.3 (27.8) | 68.7 (30.9) | 1.63 | 0.11 | 0.462 |
| % correct ordinal positions | 72.3 (35.4) | 57.3 (37.6) | 1.45 | 0.15 | 0.411 |

Numbers in the Nap and No Nap columns represent the means, with standard deviation in parentheses. We observed no group differences in explicit awareness of the motor sequences learned in Session 1 and Session 2 of Experiment 1, as measured by % correct transitions and % correct ordinal positions self-generated by the participants.

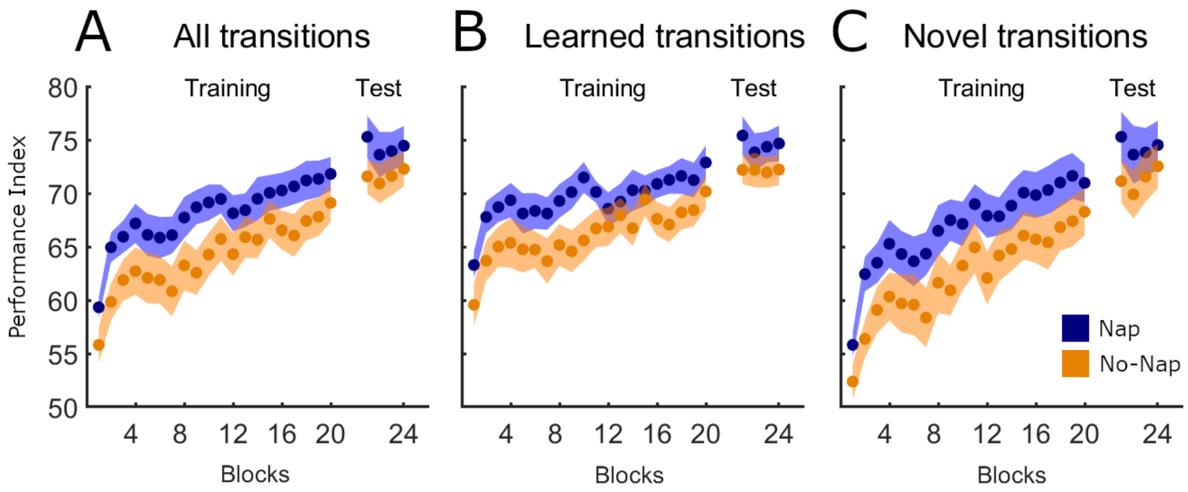


*Figure S3. Aggregate speed-accuracy measure performance index (PI) during session 2 of Experiment 1 is depicted for all (**A**), learned (**B**) and novel transitions (**C**). A higher PI index corresponds to better performance. Corresponding statistical analyses can be found in Supplementary Table 4 below. Shaded areas represent the SEM.*

*Table S4: Output of statistical analyses performed on PI in Session 2 of Experiment 1.*

| Effect | df | F | p | Partial $\eta^2$ |
|---|---|---|---|---|
| ***A. All transitions*** | | | | |
| *Training* | | | | |
| Block | 8.7,429.1 | 23.84 | <0.001* | 0.332 |
| Block x Group | 8.7,429.1 | 0.54 | 0.84 | 0.011 |
| Group | 1,48 | 3.04 | 0.09 | 0.060 |
| *Test* | | | | |
| Block | 3,138 | 1.19 | 0.32 | 0.025 |
| Block x Group | 3,138 | 0.32 | 0.81 | 0.007 |
| Group | 1,46 | 1.36 | 0.25 | 0.029 |
| ***B. Learned transitions*** | | | | |
| *Training* | | | | |
| Block | 9.2,442.2 | 9.87 | <0.001* | 0.170 |
| Block x Group | 9.2,442.2 | 0.83 | 0.59 | 0.017 |
| Group | 1,48 | 3.07 | 0.09 | 0.060 |
| *Test* | | | | |
| Block | 3,138 | 0.43 | 0.73 | 0.009 |
| Block x Group | 3,138 | 0.23 | 0.87 | 0.005 |
| Group | 1,46 | 1.56 | 0.22 | 0.033 |
| **C. *Novel transitions*** | | | | |
| *Training* | | | | |
| Block | 9.7,463.5 | 26.09 | <0.001* | 0.352 |
| Block x Group | 9.7,463.5 | 0.40 | 0.94 | 0.008 |
| Group | 1,48 | 2.38 | 0.11 | 0.054 |
| *Test* | | | | |
| Block | 3,138 | 1.55 | 0.21 | 0.033 |
| Block x Group | 3,138 | 0.62 | 0.61 | 0.013 |
| Group | 1,46 | 1.09 | 0.30 | 0.023 |
| ***D. Transition type x Group*** | | | | |
| Transition type | 1,48 | 0.23 | 0.64 | 0.005 |
| Transition type x Group | 1,48 | 0.13 | 0.72 | 0.003 |
| Group | 1,48 | 1.34 | 0.25 | 0.027 |

Output of statistical analyses on Performance Index (PI) during Session 2 of Experiment 1. Block x Group ANOVAs were computed for the training and test runs for all (**A**), learned (**B**) and novel (**C**) transitions. Additionally, a 2 (Transition type) by 2 (Group) ANOVA was also computed for performance during the test phase (**D**). These analyses correspond to those performed on the primary variables RT and Accuracy (see Figures 2-3 and Table 3 in main text) and obtained largely the same results: a significant effect of block was present for all transitions as well as for both transition subtypes, but there was no significant effect of group or group x block interaction. Performance also did not differ between the two transition types (learned and novel) nor experimental groups.

*Table S5: Correlations between sleep features (Nap group only, N=25) and performance in the sequential SRT task during Session 2 for Experiment 1.*

| | **All transitions** | **Learned transitions** | **Novel transitions** |
|---|---|---|---|
| **A. Performance Index** | | | |
| NREM duration | r=-0.24; p=0.25 | r=-0.36; p=0.08 | r=-0.12; p=0.57 |
| Spindle density | r=-0.05; p=0.81 | r=0.10; p=0.63 | r=-0.16; p=0.46 |
| Spindle amplitude | r=-0.26; p=0.20 | r=-0.19; p=0.36 | r=-0.28; p=0.18 |
| Slow wave density | r=-0.15; p=0.48 | r=-0.10; p=0.63 | r=-0.16; p=0.46 |
| Slow wave amplitude | r=-0.21; p=0.32 | r=-0.30; p=0.15 | r=-0.11; p=0.61 |
| **B. Response Time** | | | |
| NREM duration | r=0.17; p=0.41 | r=0.22; p=0.30 | r=0.14; p=0.51 |
| Spindle density | r=0.08; p=0.71 | r=-0.02; p=0.91 | r=0.15; p=0.49 |
| Spindle amplitude | r=0.19; p=0.36 | r=0.16; p=0.45 | r=0.21; p=0.33 |
| Slow wave density | r=0.10; p=0.63 | r=0.10; p=0.65 | r=0.10; p=0.63 |
| Slow wave amplitude | r=0.16; p=0.44 | r=0.21; p=0.31 | r=0.12; p=0.57 |
| **C. Accuracy** | | | |
| NREM duration | r=-0.03; p=0.88 | r=-0.19; p=0.36 | r=0.09; p=0.67 |
| Spindle density | r=-0.02; p=0.94 | r=0.02; p=0.92 | r=-0.03; p=0.88 |
| Spindle amplitude | r=-0.10; p=0.64 | r=0.01; p=0.96 | r=-0.10; p=0.62 |
| Slow wave density | r=0.08; p=0.72 | r=0.15; p=0.47 | r=-0.01; p=0.95 |
| Slow wave amplitude | r=-0.002; p=0.99 | r=-0.09; p=0.67 | r=0.06; p=0.79 |

Correlations between sleep features and sequential SRT performance in Session 2 (post-sleep), as measured by the Performance Index (**A**), Response Time (**B**) and Accuracy (**C**). For all three variables, performance was normalized by dividing the mean across the 20 training blocks of session 2 sequential SRTT by the average performance on the 4 blocks of pseudorandom SRTT completed during session 2. No significant correlations were observed. All reported values are uncorrected for multiple comparisons. Note that correlations with Performance Index were part of our pre-registered analyses, whereas those with Response Time and accuracy were considered exploratory.

*Table S8: Participant characteristics and assessments of sleep and vigilance for Experiment 2.*

| **A. Variable** | **t** | **p** | **Cohen's d** |
|---|---|---|---|
| Age | -0.84 | 0.41 | -0.221 |
| BAI score | 0.94 | 0.35 | 0.250 |
| BDI score | 0.15 | 0.88 | 0.039 |
| Handedness score | -0.04 | 0.97 | -0.010 |
| PSQI score | 0.54 | 0.59 | 0.144 |
| Daytime sleepiness score | 0.04 | 0.97 | 0.009 |
| Sleep duration, 3 nights prior to S2 | 1.76 | 0.08 | 0.471 |
| SMS duration | 1.50 | 0.14 | 0.397 |
| SMS quality | 1.03 | 0.31 | 0.273 |
| **B. SSS** | **F** | **p** | **Partial $\eta^2$** |
| Session | 2.06 | 0.16 | 0.036 |
| Session x Group | 0.54 | 0.47 | 0.010 |
| Group | 0.02 | 0.90 | <0.001 |
| **C. PVT** | **F** | **p** | **Partial $\eta^2$** |
| Session | 4.80 | 0.03* | 0.080 |
| Session x Group | 1.33 | 0.25 | 0.024 |
| Group | 0.29 | 0.59 | 0.005 |

Output of statistical analyses assessing group differences (AM-PM vs. PM-AM) in participant characteristics, sleep quality and quantity prior to the experimental sessions as well as subjective (Stanford Sleepiness Scale (SSS); Hoddes, Dement, & Zarcone, 1972) and objective (Psychomotor Vigilance Task (PVT); Dinges & Powell, 1985) assessments of vigilance. Means and SDs are provided in Table 1 of the main text. Variables in section **A** were assessed with independent samples t-tests (df = 55 for all). SSS (section **B**) and PVT (**C**) were assessed with 2 (Session) by 2 (Group) ANOVAs (df = 1,55 for all effects). No significant Group, Session or Group by Session effects were revealed. BAI = Beck's anxiety inventory [3]; BDI = Beck's depression inventory [4]; PSQI = Pittsburgh Sleep Quality Index [5], SMS = St. Mary's sleep questionnaire [6]; S2= session 2. Note that the significant effect of Session for the measure PVT was driven by overall decreased response time in Session 2 (276 ± 44 ms, compared with 286 ± 55 ms in Session 1), likely reflecting increased familiarity with the task.

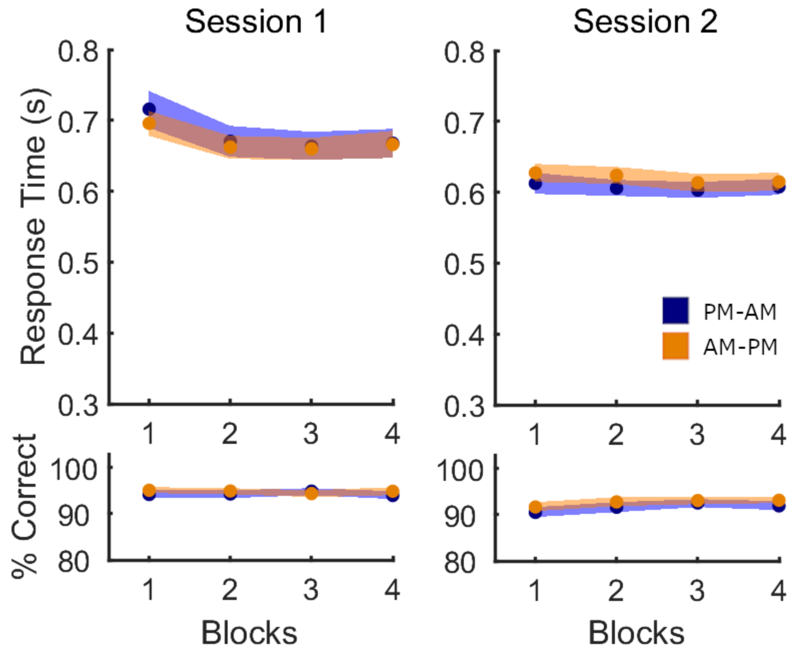


*Figure S4: Performance on the pseudo-random SRTT in Experiment 2. Mean response time (in seconds) and % correct transitions per block of sequence task are depicted for the two sessions. Output of the corresponding statistical analyses is provided in Table S9 below.*

*Table S9. Performance on pseudo-random SRT task in Experiment 2.*

| Effect | df | F | p | Partial $\eta^2$ |
|---|---|---|---|---|
| **A. Response Time (RT)** | | | | |
| *Session 1* | | | | |
| Block | 2.6,143.4 | 19.96 | <0.001* | 0.266 |
| Block x Group | 2.6,143.4 | 0.79 | 0.48 | 0.014 |
| Group | 1,55 | 0.10 | 0.76 | 0.002 |
| *Session 2* | | | | |
| Block | 3,165 | 2.21 | 0.09 | 0.039 |
| Block x Group | 3,165 | 0.50 | 0.69 | 0.009 |
| Group | 1,55 | 0.59 | 0.45 | 0.011 |
| **B. Accuracy** | | | | |
| *Session 1* | | | | |
| Block | 2.8,153.9 | 0.05 | 0.98 | 0.001 |
| Block x Group | 2.8,153.9 | 0.57 | 0.63 | 0.010 |
| Group | 1,55 | 036 | 0.55 | 0.006 |
| *Session 2* | | | | |
| Block | 3,165 | 3.38 | <0.05* | 0.058 |
| Block x Group | 3,165 | 0.16 | 0.93 | 0.003 |
| Group | 1,55 | 0.79 | 0.38 | 0.014 |
| **C. Performance Index (PI)** | | | | |
| *Session 1* | | | | |
| Block | 2.7,145.8 | 7,64 | 0.001* | 0.103 |
| Block x Group | 2.7,145.8 | 0.57 | 0.64 | 0.012 |
| Group | 1,55 | 0.20 | 0.31 | 0.021 |
| *Session 2* | | | | |
| Block | 3,165 | 4.73 | 0.003* | 0.079 |
| Block x Group | 3,165 | 0.18 | 0.91 | 0.003 |
| Group | 1,55 | 0.02 | 0.88 | <0.001 |

Output of statistical analyses assessing group differences in performance on the pseudo-random SRT task administered prior to and following the sequential SRT task in Sessions 1 and 2, respectively, reflecting general motor execution. Separate 4 (Block) by 2 (Group) ANOVAs were run per each variable (**A**: Response Time, RT; **B**: Accuracy) and each Session. The presence of a significant effect of Block during Session 1 for RT and PI indicates a general increase in performance, presumably due to task familiarization. The significant effect of *block* on accuracy in Session 2 was caused by a marginal decrease of an overall very high accuracy during the first block (average 91% accuracy, compared with 92% in the other three blocks. No main effect of group or block x group interaction were observed for any measure, in either session, demonstrating that general motor execution did not differ between experimental groups. Df = degrees of freedom.

*Table S10. Performance on the generation task in Experiment 2.*

| **Variable** | **AM-PM** | **PM-AM** | **t(55)** | **p** | **Cohen's d** |
|---|---|---|---|---|---|
| *Session 1* | | | | | |
| % correct transitions | 13.3 (24.8) | 7.8 (11.7) | 1.08 | 0.28 | 0.287 |
| % correct ordinal positions | 24.0 (23.1) | 19.2 (12.3) | 0.97 | 0.34 | 0.257 |
| *Session 2* | | | | | |
| % correct transitions | 43.5 (40.6) | 52.5 (40.4) | -0.84 | 0.41 | -0.223 |
| % correct ordinal positions | 48.3 (37.8) | 59.4 (35.0) | -1.15 | 0.26 | -0.304 |

Numbers in the AM-PM and PM-AM columns represent the means, with standard deviation in parentheses.

We observed no group differences in explicit awareness of the motor sequences learned in Session 1 and Session 2 of Experiment 1, as measured by % correct transitions and % correct ordinal positions self-generated by the participants.
